# Visual and vestibular reweighting after cyber- and space-sickness

**DOI:** 10.1101/2025.03.12.642770

**Authors:** Tess Bonnard, Emilie Doat, Jean-René Cazalets, Dominique Guehl, Etienne Guillaud

## Abstract

Sensory conflicts are widely recognized as the primary drivers of motion sickness (MS), though the underlying integrative processes remain poorly understood. This study investigated sensory reweighting that follows exposure to different sensory conflict paradigms. First, visual and vestibular reflexes were assessed before and after a visuo-vestibular conflict induced by purely visual stimulation in virtual reality. Second, visual and vestibular integration were evaluated before and after an otolith-canal conflict induced by gravitational changes in parabolic flight. Semi-circular canal integration was measured via the vestibuloocular reflex (VOR) suppression task, while visual weighting was assessed through optokinetic nystagmus (OKN). Our findings revealed that different sensory conflict paradigms elicit distinct sensory reweighting processes. Visuo-vestibular conflict resulted in a decreased VOR response, whereas vestibulo-vestibular conflict mainly led to a reduction in OKN following parabolic flight. Sensory down-weighting occured in the modality that did not detected displacement, likely perceived as the less reliable input, regardless of its accuracy. Additionally, visual sensitivity emerged as a potential predictor of cybersickness, while vestibular sensitivity seemed to influence MS severity in parabolic flight. Our data suggest that the sensitivity of the most stimulated sensory modality during a given conflict may determine an individual’s susceptibility to MS.

**KEY POINTS:**

- Sensory reweighting occur through brief and specific exposure to motion sickness.
- Adaptive reweighting is modulated by the nature of the motion sickness exposure, with distinct effects observed between space-sickness and Earth-like motion sickness cues.
- Motionless cues are consistently downweighted, regardless of their accuracy.
- Motion sickness intensity depends on individual’s sensitivity to the stimulated sensory sources, which varies across provocative sensory environments.

## INTRODUCTION

Motion sickness (MS) affects a substantial portion of the population, with symptoms ranging from mild discomfort (e.g., nausea, pallor, cold sweats) to more severe impairments, including cognitive and physical performance deficits (Golding, 2016 ; Zhang, 2016). These symptoms can have critical implications, potentially hindering astronauts’ ability to perform highly skilled tasks during the initial days of space travel (Heer, 2006) or reducing alertness and raising safety concerns for car passengers (Pereira, 2024). Current treatments, such as pharmacological solutions, often carry side effects that reduce alertness, increase imbalance in already unstable environment, or induce drowsiness, which are all incompatible with tasks requiring focus and precision (Weerts, 2014; Bestaven, 2016). Despite its prevalence, the integrative mechanisms underlying MS remain poorly understood and largely theoretical, with limited identification of effective predictive factors.

The MS symptoms might arise from disruptions in the brain’s ability to integrate sensory information about motion and spatial orientation. Sensory signals from multiple modalities converge in cortical and subcortical structures, where they are integrated and weighted based on their consistency with other sensory inputs (Spence, 2004; Choi, 2023). While the human sensory systems has evolved over millions of years, the integration of new sensory combinations has been introduced over the last century by the rapid emergence of modern technologies (car, plane, virtual reality, …). These environments often create sensory discrepancies that human brains are not yet adapted to resolve. The sensory conflict theory, one of the most widely accepted frameworks, posits that MS arises from contradictory sensory inputs (Guedry, 1998; Reason & Brand, 1975 ; Bos, 2011 ; Irmak, 2023). When sensory systems provide conflicting information about motion or orientation, the CNS struggles to reconcile these signals, leading to discomfort and other symptoms.

MS can be categorized based on the sensory modalities that trigger it. Broadly, two main types can be distinguished : visuo-vestibular (inter-sensory) conflict and vestibulo-vestibular (intra-sensory) conflict (Reason, 1978 ; Gallagher & Ferrè, 2018). Visuo-vestibular conflict arises from incoherences between visual and vestibular inputs and is commonly experienced during various modes of transportation, such as reading in a moving car or traveling on a windowless boat. It is also prevalent in emerging virtual reality (VR) technologies. In VR, users are immersed in a full-field artificial visual environment that induces a sensation of vection (self-motion perception), despite their physical body remaining largely static, seated, or exhibiting restricted movement. This sensory conflict between visual motion perception and the absence of corresponding bodily movement results in cybersickness (Kim, 2005), which affects approximately 60 to 95% of VR users (Cobb, 1999 ; Caserman, 2021). In contrast, vestibulo-vestibular conflict does not stem from contradictions between different sensory systems but rather from inconsistencies within the vestibular system itself. This phenomenon occurs in environments with altered gravitational conditions, such as space-flight or parabolic flights, where individuals are suddenly exposed to weightlessness. Humans, having evolved under Earth’s gravity, are accustomed to a relatively stable gravitoinertial vector as detected by the otolith organs, with only minor variations. This otolithic input is typically congruent with semicircular canal detection of head rotations, forming the basis of vestibular processing that the brain has adapted to over evolutionary time (Bertolini, 2021 ; Merfeld, 2005a, 2005b). However, in highly altered gravitational environments, the otoliths provide unexpected signals, while the semicircular canals continue detecting angular accelerations in the usual manner. This novel sensory combination leads to an otolithocanalar conflict which has been proposed as the origin of space motion sickness (SMS) (Graybiel, 1975; Graybiel & Lackner, 1977). SMS affects approximately two-thirds of astronauts during their initial days in space and one-thirds of medicated flyers during parabolic flights (Lackner & DiZio, 2006 ; Golding, 2017). Notably, prolonged exposure to weightlessness leads to otolith tilt-translation reinterpretation (Parker, 1985), a phenomenon in which otolithic stimulation become perceived more as translation than tilt, mirroring the conditions experienced in space. This reinterpretation highlights the importance of adaptative internal models in sensory integration and serves as the foundation for another approach to explaining MS : the Neural Mismatch Theory. This theory suggests that MS arises from discrepancies between current sensory inputs and prior expectations based on past experiences (Reason, 1978). Ultimately, both sensory conflict and neural mismatch theories emphasize the central role of sensory integration in the development of MS.

Multisensory integration relies on the convergence of various sensory inputs generated by a single stimulus, using a weighted linear combination of individual perceptual estimates (Ernst & Banks, 2002). It has been proposed that this process serves to reduce perceptual uncertainty about the stimulus (Knill & Pouget, 2004). When one sensory signal is incongruent with the majority of other signals, it introduces greater uncertainty into the integrative process, compromising the accuracy of perception. To address this, the integration mechanism tends to suppress the source of uncertainty, ensuring a more coherent and reliable final perception. The Maximum Likelihood Estimation (MLE) model incorporates these principles, essentially functioning on the idea that “the most reliable input will be weighted more heavily than others in the final integration” (Ernst & Bülthoff, 2004). According to the MLE model, space sickness resulting from otolith-canal conflict could diminish the perceived reliability of vestibular input, leading to a downweighting of vestibular cues. Conversely, intense visual stimulation in a stationary participant could compromise the credibility of visual input, reducing the weight assigned to visual motion detection.

Our study aims to deepen the understanding of sensory weighting modulation by examining how these two distinct sensory conflict paradigms impact low-order reflexes. Specifically, we investigated whether sensory conflicts induced VR and parabolic flights lead to different sensory weighting strategies or whether MS, regardless of its origin, produces similar effects on sensory integration. We assessed the effects of a single short-term exposure (10–20 minutes) to a provocative situation, known to be sufficient to trigger MS symptoms, on vestibulo-ocular reflexes (VOR) and optokinetic reflex (OKN). Additionally, we explored potential relationships between sensory processing mechanisms and individual susceptibility to MS severity. We hypothesized that the sensory modality most affected by stimulation during conflict would be perceived as less reliable, leading to a post-exposure reduction in sensitivity to motion cues from that modality. Specifically, we expected a decrease in OKN following the virtual reality session and impaired VOR performance after the parabolic flight session.

## MATERIAL AND METHODS

### Participants

29 healthy participants in total were recruted for the project (27 ± 7 years ; 14 females, 15 males). Among the 23 participants participated to parabolic flight (PF) protocol (28 ± 7 years ; 10 females, 13 males), 21 of them also performed the virtual reality (VR) protocol in the laboratory (27 ± 7 years ; 9 females, 12 males). 6 participants did only the VR protocol (22 ± 2 years ; 4 females, 2 males). For participants enrolled in both VR and PF protocols, half of them did the VR protocol one month before the PF protocol and the other half performed the VR protocol one month after the PF protocol, to prevent protocol order bias. No specific sensory testing was performed prior to the experiments. None of the participants reported any health issues or previous medical history which could have impacted their sensory functions. For participants selected for the PF protocol, a medical form was filled and approved by an aerospatial physician, based on the analysis of participant’s effort electrocardiogram made by a cardiologist. Before taking part in the present study, all subjects provided a signed consent form after a cooling-off period, and after inclusion visit for PF protocol. Ethical approval for this study was granted by the national Ethics Committee « Comité de protection des personnes Ile de France II » n°2023-A02145-40 and all experiments were conducted according to Helsinki Agreement.

### Sensori-motor tests

Vestibulo-ocular reflexes and optokinetic nystagmus were evaluated before and after exposure to VR and parabolic flights (figure 1). Posturographic tests were performed in laboratory, before and after VR exposure.

**Figure 1.**
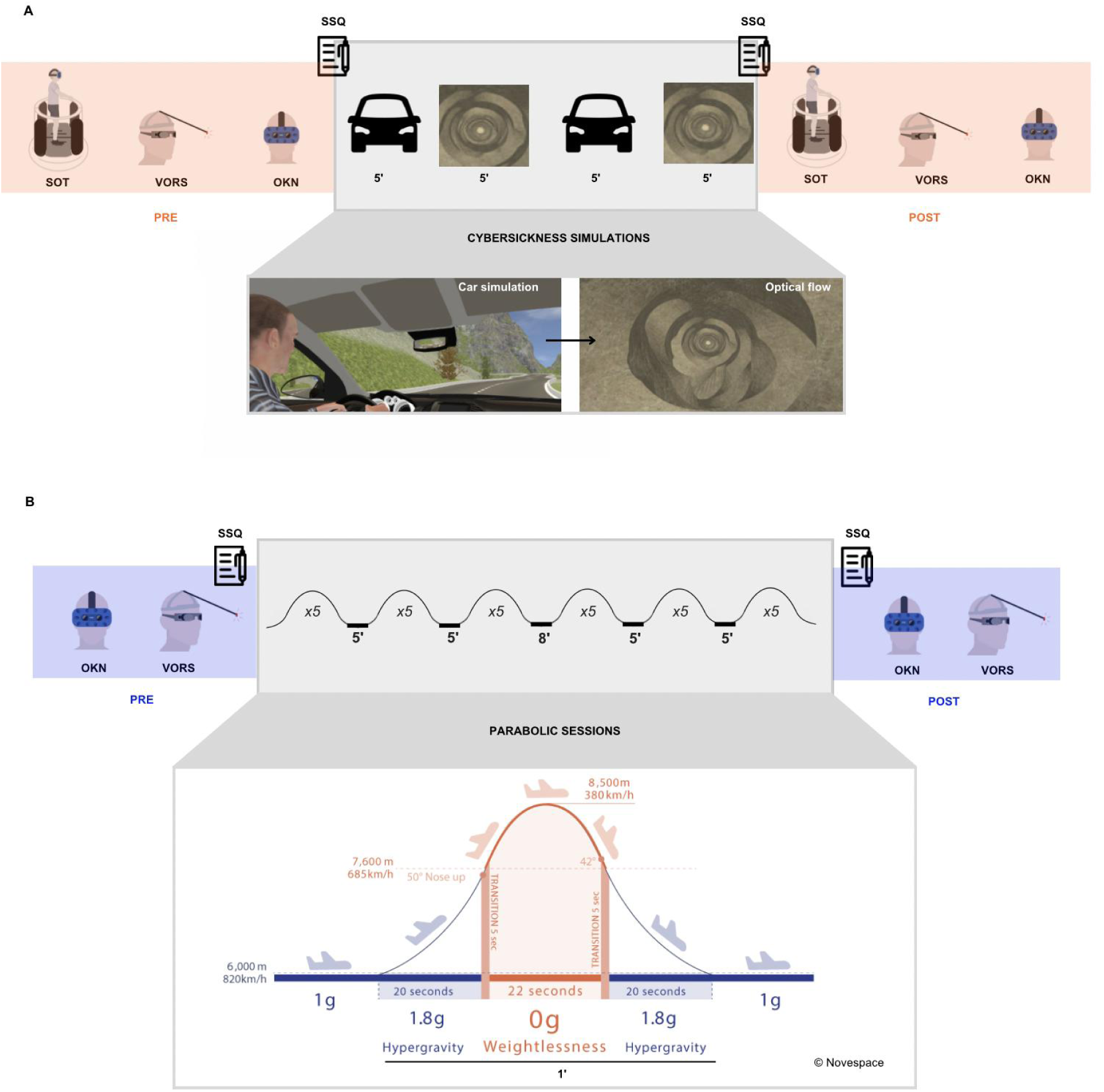
VR protocol timeline (A) and PF protocol timeline (B). The VR protocol lasted approximately 2 hours, whereas the PF protocol extended to about 3 hours. Times are presented in minutes.

**Figure 2.**
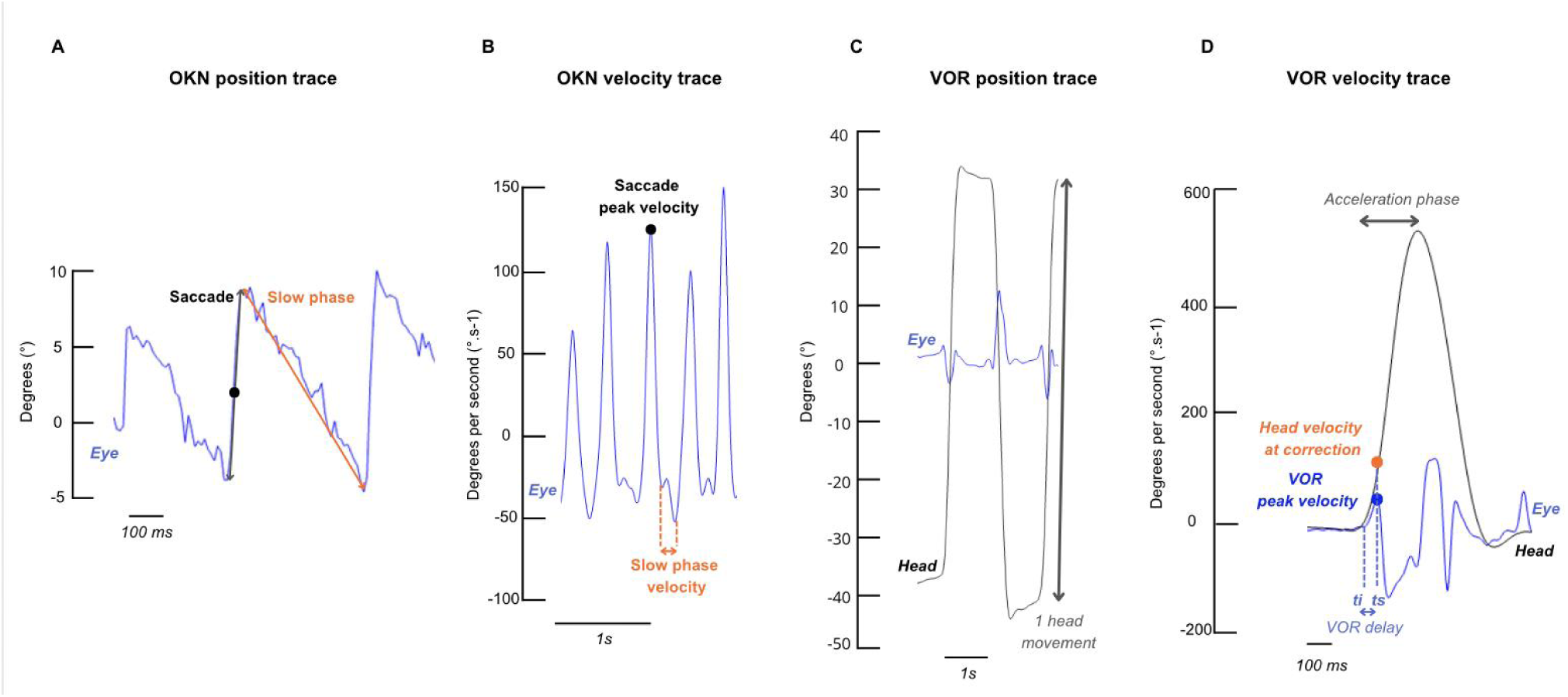
Eye position and velocity traces during OKN (A-B) and VORS (C-D) testing. For OKN (A), eye position (blue) is divided into two phases : the slow phase (orange double-sided arrow) where the eye tracks the moving visual stimulus, and the corrective saccade (grey double-sided arrow) that recaptures the visual input on the retina. Saccades between slow phases are clearly identified by velocity peaks (B). For VOR (C), both eye (blue) and head (black) movements are illustrated. Head movements are fast, large, and involve a 1 second break between directional changes. The eye’s corrective reflex, which opposes head movement, occurs during the head acceleration phase and represents the residual VOR response before suppression (D). Any eye movements occurring after the head acceleration phase can be caused by different strategies such as smooth pursuit or fixation. ti = time of head movement initiation ; ts = time of VORS

***Optokinetic nysgtamus reflex (OKN)*** was tested using a bilateral eye-tracking system implemented inside a VR headset (HTC Vive ProEye). White dots on a black blackground moving at 30°/s from right-to-left, left-to-right, top-to-bottom and bottom-to-top respectively was generated by the OKN software from Virtualis (Montpellier, France) (Figure 3A). The same presentation order was kept pre- and post-exposure, and for each participant. Each direction recording lasted 30s. Participants were asked to look straight forward before the start of stimulation and to let their eyes move freely without controlling them.

**Figure 3.**
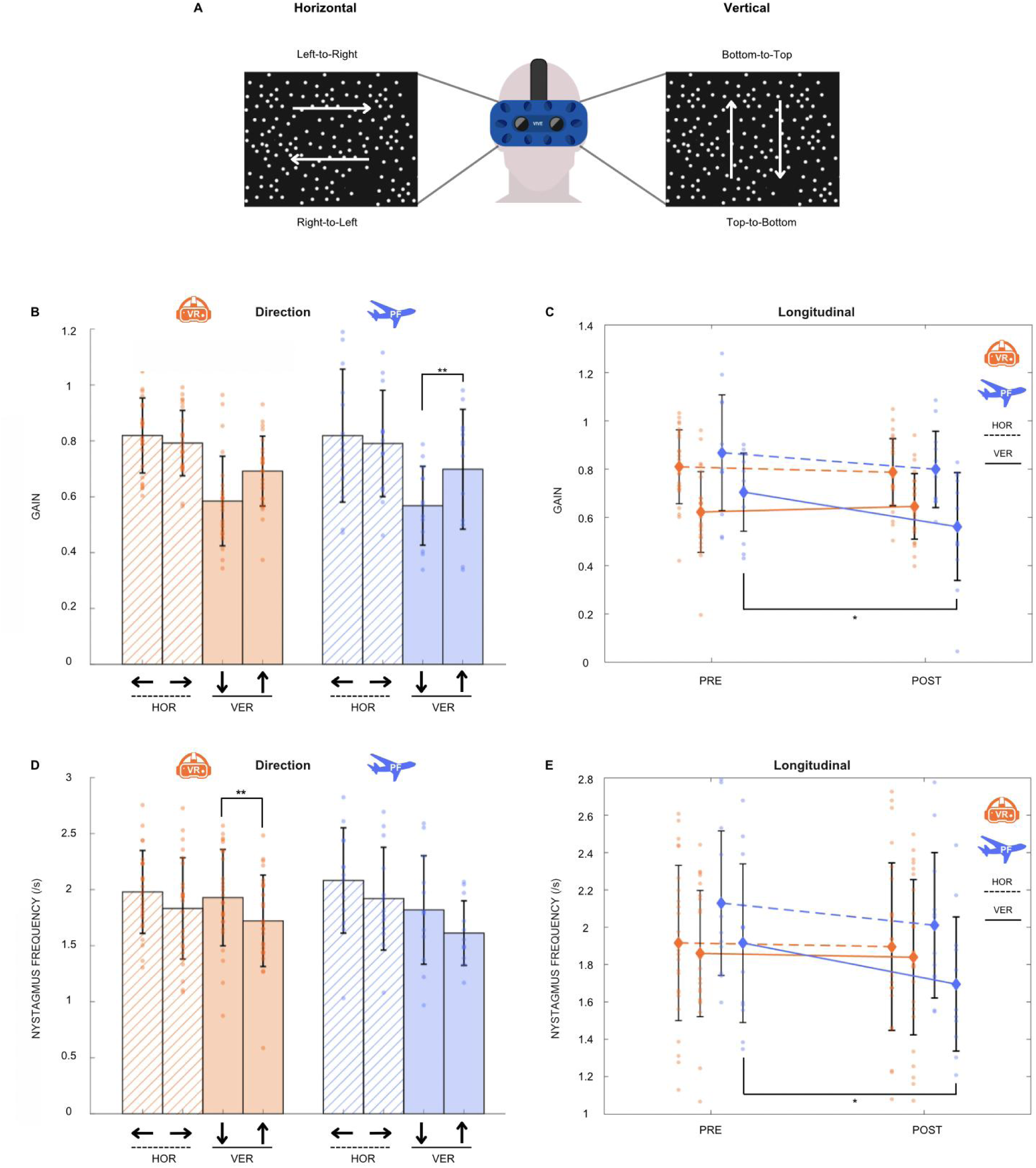
OKN results from VR and PF protocols. Directions are referred to as Right-to-Left, Left- to-Right, Top-to-Bottom, Bottom-to-Top and are described in panel A. Gain (B-C) and nystagmus frequency (D-E) are shown for both paradigms (Virtual reality = VR & Parabolic flight = PF) and orientations (HOR = Horizontal & VER = Vertical). Repeated-measure ANOVA, Fisher’s LSD post-hoc, VR : n = 25, PF : n = 13, * : p <0.05 ; ** : p<0.01.

***Vestibulo-ocular reflex suppression (VORS)*** was evaluated by recording the movement of the right eye pupil using an infrared camera on eye-tracking googles equipped with an integrated gyroscope to record head movements (Otometrics ICS Impulse, Natus). To avoid visual disturbance, occulting lenses were fitted on googles, allowing specific far-red and infrared wavelengths to pass through. A lab-made helmet composed of a carbon stem ended with a red LED (45 cm in front of the participant) was placed and tighten on the participant head. The stem was adjusted to place the red target in the middle of the visual field of the participant (Figure 4A). Participants were asked by the experimenter to voluntary move their head quickly in the horizontal or in the vertical orientation and to make 1s break before starting any movement in the other direction (e.g., from right-to-left to left-to-right). The task required maintaining stable eye fixation during head rotation, with the gaze fixed on the red LED of reference that rotated along with the head. A horizontal trial was performed first and vertical second, with each lasting between 30 to 40s, with 20 head movements realized per trial. This task was repeated 3 times.

**Figure 4.**
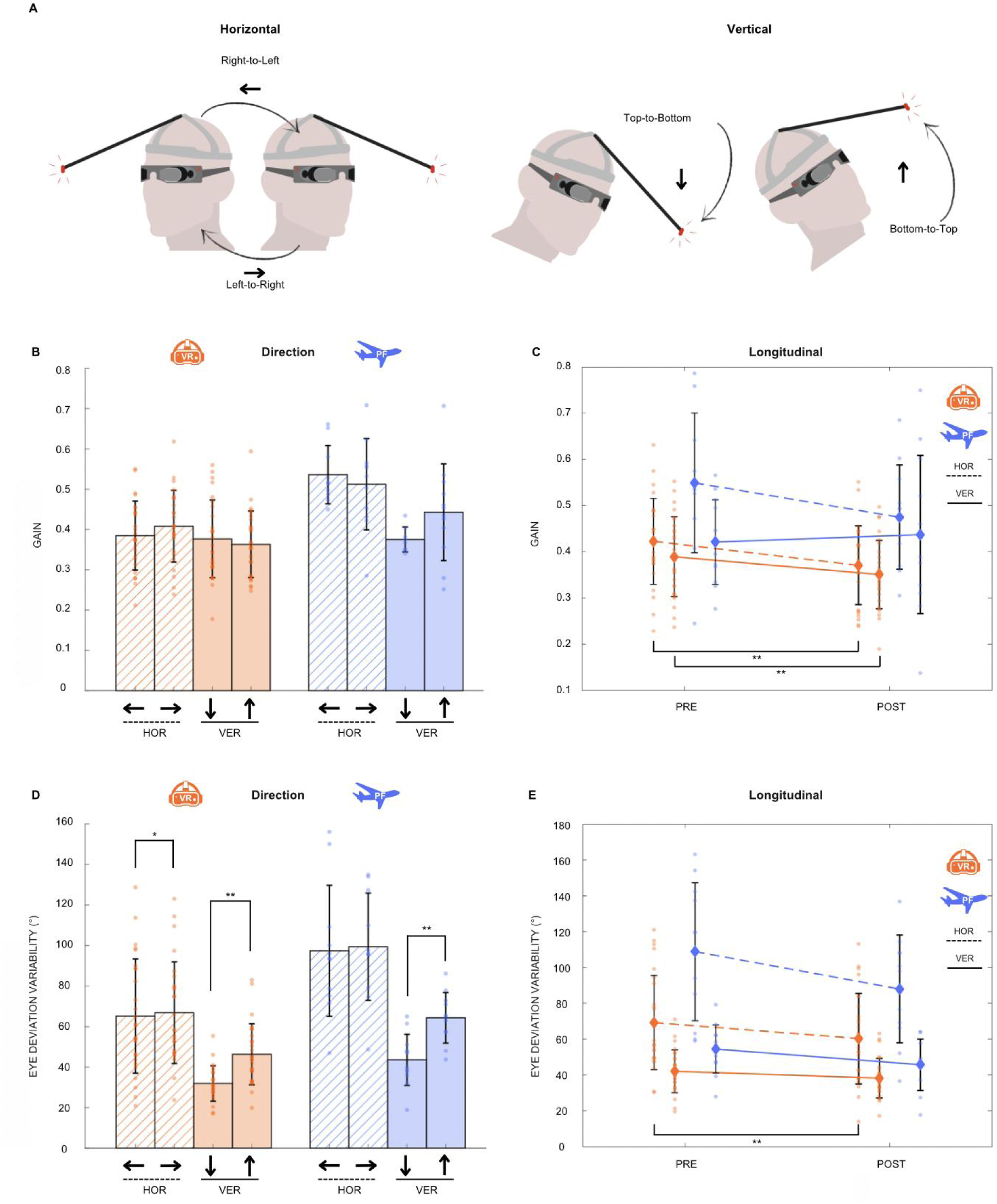
VORS results from VR (B-C) and PF (D-E) protocols. Directions are referred to as Right-to-Left, Left-to-Right, Top-to-Bottom, Bottom-to-Top and are described in panel A. Gain-at-correction (B-C) and eye deviation variability (D-E) are shown for both paradigms (Virtual reality = VR & Parabolic flight = PF) and orientations (HOR = Horizontal, VER = Vertical). Repeated-measure ANOVA, Fisher’s LSD post-hoc, VR : n = 27, PF : n = 11-12, * : p <0.05 ; ** : p<0.01.

**Figure 5.**
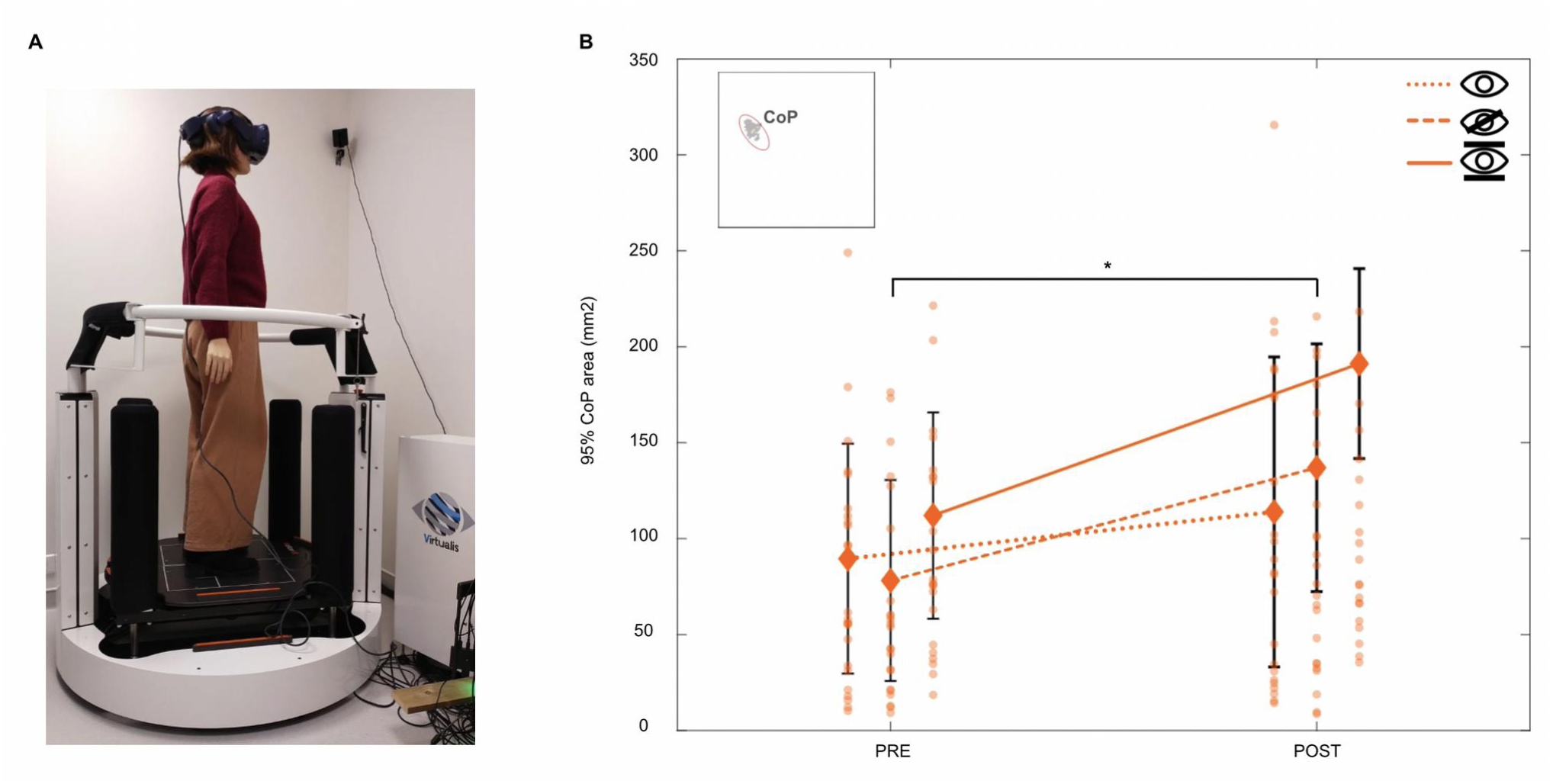
SOT results before and after VR sensory conflict. Panel A illustrates the SOT setup (the person in the photograph is one of the authors). The feet center-of-pressure (CoP) with the three experimental conditions are displayed as legends in Panel B. They are presented as followed : (dotted line) normal visual input, (dashed line) vision deprivation, and (full line) fixed visual environment despite head movements. The platform measuring the CoP remained stable during all these conditions The 95% area represents the elliptical region (red ellipse, Panel B) encompassing 95% of the CoP displacement during the trials. Statistical significance was assessed using non-parametric Wilcoxon signed-rank test (n = 27). Significance levels are indicated as follows : * p < 0.05, ** p < 0.01.

Posturographic Test was conducted using the Virtualis platform (MotionVR) and its standardized ***sensory organization test (SOT)*** protocol (Figure 6A). The protocol included three conditions, each comprising three 20-second trials. The visual conditions were displayed as followed : (1) normal vision dynamically adjusted to head movements, (2) vision deprivation (black screen), and (3) fixed vision, where the visual scene remained unchanged despite head movements. These visual conditions were assessed while the platform was static. Participants were instructed to maintain a forward gaze and remain as still and stable as possible throughout each trial. Feet center-of-pressure (CoP) was measured by force detectors integrated into the platform to observe posturographic variations (Figure 6A).

**Figure 6.**
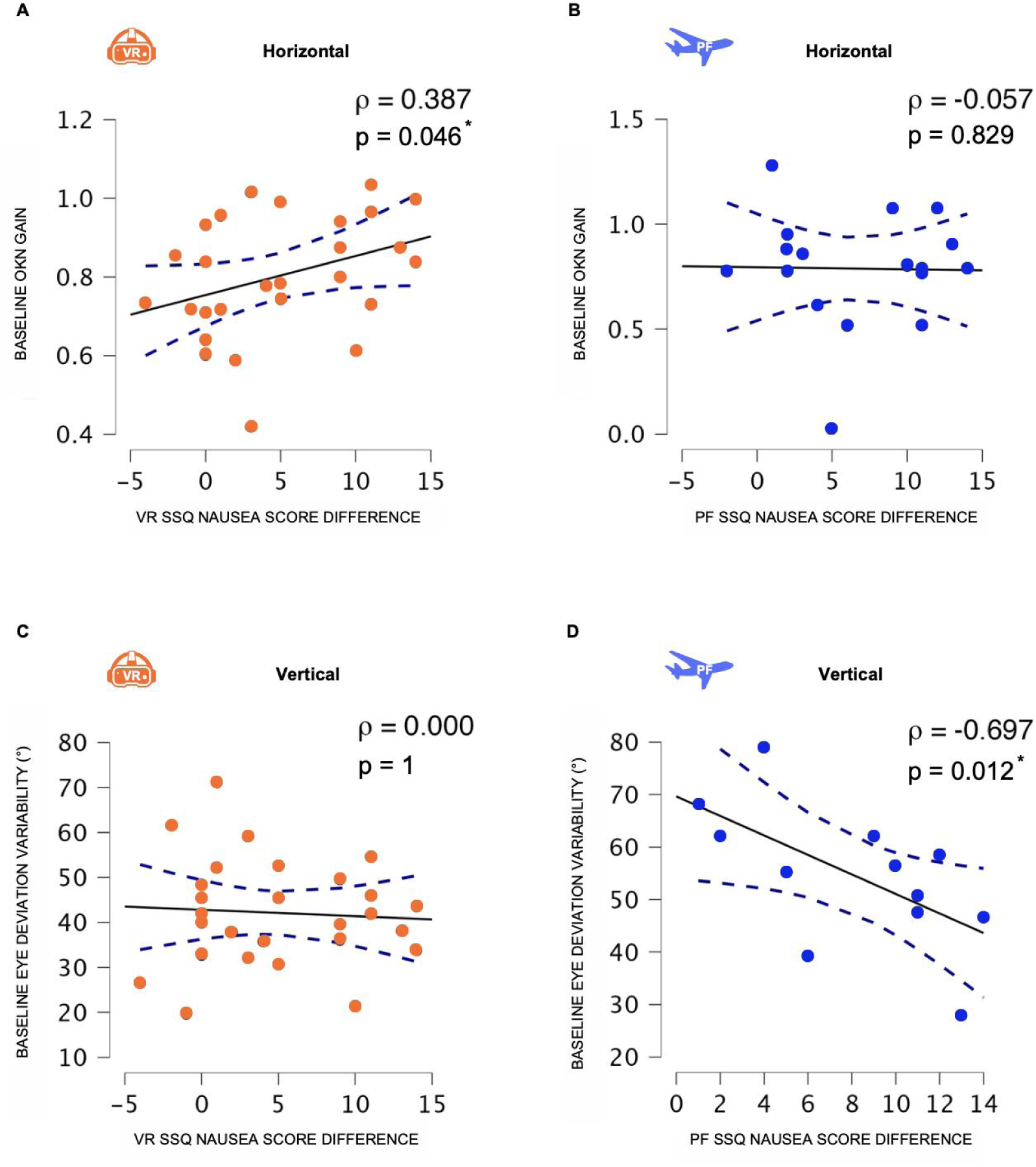
Correlations between sensory performance and motion sickness severity. Correlations between SSQ nausea score differences and OKN gain (A-B), and between SSQ nausea score differences and baseline eye deviation variability (C-D) are presented. Baseline eye deviation variability in parabolic flight refers to pre-flight values measured before take-off. Spearman correlations. VR : n = 27, PF : n = 12-17, ρ < 0.3 low, ρ <0.5 moderate, ρ > 0.5 high.

*Vestibular-evoked myogenic potentials (VEMP)* were assessed during both protocols. This test relied on the recording of otolithic-driven muscle reflexes elicited by vestibular stimulation using tone-burst auditory stimulation. However, no significant longitudinal effects were observed in either, and thus these results were not included in this paper.

### Virtual reality protocol

Sensori-motor tests were conducted in a specific order, reproduced for each participant, starting with the most comfortable test and finishing with the most provocative one. They first performed the balance SOT, followed by VORS and finished with OKN (Figure 1 A). Those experimental results were accounted as baseline responses. Participants were then placed upright on a platform with a virtual reality headset for the sensory conflict procedure (MotionVR, Virtualis). They watched two different simulations : one as a front side car passenger on a mountain road, and the other inside a tunnel which rotated in a random orientation every 10s and switch from forward to backward motion every 8s. Each simulation lasted 5 min and were repeated twice. Continuous dialogue was kept between the participant and the experimenter to follow the participant subjective state. Sensory conflict procedure was stopped when participants reported a moderate nausea or inco mfort (Motion Sickness Severety Scale = 4 ; Czeiler, 2023) or when they requested it. Participants without symptoms remained on the simulation for a total of 20 min. Sensori-motor evaluations were performed post-conflict in the same order (SOT>VORS>OKN) (Figure 1A). SSQ questionnaires were filled right before sensory conflict, right after and at the end of the protocol.

### Parabolic flight protocol

Parabolic flights were conducted during the VP177 and VP181 parabolic flight campaigns of the Centre National d’Etudes Spatiales (CNES) aboard the Airbus A310 Zero G, operated by Novespace in Mérignac (Bordeaux, France). Each parabolic flight consisted of 31 parabolic maneuvers, each following a sequence of altered gravity phases : hypergravity (1.8g), microgravity (0g), and a return to hypergravity (1.8g), interspersed with periods of normogravity during steady flight (Figure 1B). Each parabola lasted one minute, with a two-minute interval between successive parabolas. Five-minute breaks were scheduled after every five parabolas, with an extended eight-minute break at the midpoint of the flight (Figure 1B). The total flight duration ranged from 2.5 to 3 hours, encompassing takeoff, transit to the flight zone, the parabolic session, return to the airport, and landing. To prevent MS, scopolamine is commonly administered pre-flight, as it is known to attenuate vestibular integration (Weerts, 2015; Bestaven, 2016). However, in this study, no medication was given before the flight to preserve the integrity of vestibular function and maintain participant alertness.

Before takeoff, while the aircraft was stationary on the ground, participants underwent baseline measurements of OKN (four directions, 30 seconds per direction, 30 °/s) and VORS (horizontal and vertical, each repeated twice, 30 seconds per trial). Additionally, VORS was performed during approximately 10 parabolas spread throughout the parabolic session (between the 5th and 25th parabola). However, the goggle-mounted gyroscope produced incoherent kinematic data when gravity was altered, rendering the in flight test unusable. After the final parabola (during the steady flight before landing), OKN was tested again, whereas VORS was assessed after landing on ground. A schematic timeline of the experimental protocol is presented in Figure 1B. Participants were allowed to withdraw from in-flight testing or take breaks as needed. Those who were unable to continue the flight were excluded from further testing and, depending on the remaining flight duration, could receive a scopolamine injection to alleviate symptoms.

### Motion sickness evaluation

MS was evaluated by two main methods : subjective and objective measures. Subjective sensations and feelings were reported using the Simulator Sickness Questionnaire (SSQ) (Kennedy, 1993) before any experience at the beginning of the VR protocol, right after the virtual sensory conflict and at the end of all experiences. For the PF protocol, SSQ was filled before taking-off and right after the end of the flight.

### Data analysis

All analysis were performed using Matlab R2024b (The Mathworks, Natick, USA). For OKN, gaze position were low-pass filtered using a fourth-order Butterworth filter with a cutoff frequency of 5 Hz (Figure 2A). Saccades were identified algorithmically by detecting velocity peaks exceeding a threshold of 100°/s. The onset and offset of each saccade were determined by locating the first velocity inversions occurring before and after the velocity peak, respectively. All detected saccades where visually inspected after algorithm run, and mismatches were manually corrected. Eye displacements between the end of the (n-1) saccade and the start of the n-th saccade, characterized by lower velocity in the opposite direction, were considered slow phases (red arrow, Figure 2B). Saccadic and slow phase durations and amplitudes were computed. OKN gain was calculated by dividing the eye slow-phase velocity by the visual stimulus velocity (30°.s^-1^). Nystagmus frequency was determined by dividing the total number of eye slow-phases by the trial duration.

For VORS, vertical and horizontal eye positions in degrees and head yaw, pitch and roll (Figure 2C) were extracted, low-pass filtered (Butterworth, fourth-order, 20 Hz for eyes and 5 Hz for head) and differentiated to calculate velocities. Head rotations were identified when head velocity exceed 60°.s^-1^, and the onset and offset of head rotation were determined by locating the first velocity inversions occurring before and after the velocity peak, respectively. Any remaining mismatches, after algorithm detection, were visually inspected, and manually corrected if needed. The head acceleration phase was defined as the first part of head movement, between onset and velocity peak (Figure 2D). In the velocity plot, eye and head traces in the same direction indicate that the eyes are moving in the opposite direction compared to the head (Figure 2D). The task required maintaining a stable gaze on a fixation point that rotated with the head, keeping the eyes fixed relative to the head. We calculated the Eye Deviation Variability (EDV), defined as the standard deviation of eye position during the head acceleration phase. The correction point was defined as the time at which the eye movement transitioned in the direction of the head, marking the onset of VOR suppression (Figure 2D). VOR gain was calculated as the ratio between the eye peak velocity during the remaining VOR and the head velocity at the point of correction. VOR delay was defined as the time interval between the onset of the VOR (*ti*, Figure 2D), marked by the eye moving in the direction opposite to the head, and the correction point (*ts*, Figure 2D).

### Statistical analyses

Statistical analyses were conducted using MATLAB for repeated-measures ANOVA and JASP for correlation and Wilcoxon signed-rank tests. Pre-post effects and directional effects were assessed separately for horizontal and vertical data. For both on-ground and in-flight results, we performed repeated-measures ANOVA with Least Significant Difference (LSD) Fisher’s correction. Prior to running the ANOVA, Levene’s test was used to verify homogeneity of variance, which was satisfied in all analyses (p > 0.05). Mauchly ’s test of sphericity was also conducted, and if the assumption was violated (p < 0.05), the Greenhouse-Geisser correction was applied when epsilon (ε) was < 0.75, whereas the Huynh-Feldt correction was used for ε ≥ 0.75. Effect sizes for ANOVA results were reported as partial eta squared (η²), calculated as the ratio of the effect sum of squares to the sum of the effect and error sum of squares. Correlation analyses were conducted using Spearman’s ρ. SOT pre-post effects were tested using non-parametric Wilcoxon signedrank test, based on Shapiro-Wilk normality test results. Effect size was reported as matched rank biserial correlation (r) value. Results are presented as mean ± standard deviation (SD) unless specified.

## RESULTS

### Optokinetic Nystagmus

In the VR protocol, OKN exhibited horizontal and vertical amplitudes of 7.9° ± 3.7° and 7.1° ± 3.7°, respectively. The duration was 322 ms ± 117 ms (horizontal) and 333 ms ± 138 ms (vertical). Horizontal eye slow-phase velocity was 24°.s⁻¹ ± 4.6°.s⁻¹, and vertical velocity was 19.8°.s⁻¹ ± 5.8°.s⁻¹. Nystagmus frequency was approximately 1.9 ± 0.5 Hz in both orientations.

In the PF protocol, horizontal and vertical amplitudes were 7.6° ± 3.9° and 7.3° ± 3.9°, respectively. Duration was 294 ms ± 102 ms (horizontal) and 367 ms ± 128 ms (vertical). Horizontal eye slow-phase velocity was 24.6°.s⁻¹ ± 6.4°.s⁻¹, and vertical velocity was 20.3°.s⁻¹ ± 5.7°.s⁻¹. Nystagmus frequency was around 2.0 ± 0.5 Hz (horizontal) and 1.7 ± 0.5 Hz (vertical).

No significant longitudinal effects were observed for OKN gain during the VR protocol in horizontal (PRE = 0.80 ± 0.18, POST = 0.79 ± 0.16, F(1,24) = 0.5, p = 0.483 ) and in vertical (PRE = 0.64 ± 0.21, POST = 0.67 ± 0.20, F(1,25) = 1.2, p = 0.282) (Figure 3C). In contrast, parabolic flight primarily affected vertical OKN responses. A significant pre-post decrease in vertical OKN gain was observed post-flight (PRE = 0.72 ± 0.19, POST = 0.59 ± 0.23, F(1,12) = 7.8, p = 0.0164, η² = 0.39) (Figure 3C). A similar decreasing trend was noted for horizontal OKN gain post-flight, though it did not reach significance (PRE = 0.80 ± 0.18, POST = 0.78 ± 0.16, F(1,12) = 2.5, p = 0.139) (Figure 3C). In the parabolic flight experiment, a directional effect was observed with vertical OKN gain significantly higher during bottom-to-top movements (T-to-B = 0.58 ± 0.18, B-to-T = 0.72 ± 0.23, F(1,12) = 15.8, p = 0.00186, η² = 0.57) (Figure 3B). Similar but not significant effect was observed in the VR experiment (T-to-B = 0.60 ± 0.24, B-to-T = 0.67 ± 0.16, F(1,25) = 1.25, p = 0.273) (Figure 3B). No significant interaction between PRE-POST and directional effects was found in either protocol.

Regarding nystagmus frequency (Figure 3, lower panel), no significant longitudinal effects were observed in the VR protocol (Figure 3E). Specifically, there was no change in horizontal nystagmus frequency (PRE = 1.92 ± 0.54, POST = 1.90 ± 0.53 ; F(1,24) = 0.004, p = 0.949) or vertical nystagmus frequency (PRE = 1.82 ± 0.48, POST = 1.84 ± 0.46 ; F(1,25) = 0.18, p = 0.674). In contrast, in the PF protocol, a post-flight decrease in nystagmus frequency was observed, which was significant in the vertical orientation (Figure 3E, PRE = 1.83 ± 0.46, POST = 1.69 ± 0.51 ; F(1,12) = 5.7, p = 0.0346, η² = 0.32) but not significant in the horizontal direction (PRE = 2.13 ± 0.47, POST = 2.01 ± 0.58 ; F(1,12) = 2.7, p = 0.127). A directional effect was also observed, with a higher nystagmus frequency during top-to-bottom vertical movements which was significant in VR protocols (T-to-B = 1.92 ± 0.48, B-to-T = 1.73 ± 0.44, F(1,25) = 8.6, p = 0.00714, η² = 0.26) but not significant in PF protocol (T-to-B = 1.80 ± 0.54, B-to-T = 1.66 ± 0.44, F(1,12) = 2.1, p = 0.177) (Figure 3D).

Additional OKN variables showed no significant pre-post effects in either orientation during the VR. Eye slow phase amplitude remained unchanged post-exposure to VR in horizontal (PRE = 7.88 ± 3.92, POST = 7.94 ± 3.48, F(1,24) = 0.15, p = 0.704) and in vertical orientations (PRE = 6.94 ± 3.56, POST = 7.31 ± 3.92, F(1,25) = 0.66, p = 0.425). Similarly, eye slow-phase duration was stable in this protocol both in horizontal (PRE = 0.55± 1.66, POST = 0.32 ± 0.12 , F(1,24) = 0.94, p = 0.341) and in vertical orientations (PRE = 0.64 ± 2.33, POST = 0.35 ± 0.15, F(1,25) = 0.88, p = 0.357). In PF protocol, eye slow-phase amplitude was not influence by parabolic sessions in horizontal (PRE = 8.34 ± 4.57, POST = 6.89 ± 2.97, F(1,12) = 2.65, p = 0.129) and vertical orientations (PRE = 7.91 ± 4.56, POST = 6.67 ± 3.06, F(1,12) = 1.81, p = 0.203). Likewise, eye slow-phase duration was constant post-flight both in horizontally (PRE = 0.28 ± 0.11, POST = 0.51 ± 0.79, F(1,12) = 1.23, p = 0.290) and vertically (PRE = 0.36 ± 0.13, POST = 0.52 ± 0.58, F(1,12) = 1.69, p = 0.217).

In PF, vertical eye slow-phase amplitude was significantly higher during bottom-to-top movements (T-to-B = 5.95 ± 3.70, B-to-T = 8.39 ± 3.79, F(1,12) = 22, p < 0.001, η² = 0.65) whereas slow-phase duration remain stable (T-to-B = 0.43 ± 0.51, B-to-T = 0.44 ± 0.34, F(1,12) = 0.35, p = 0.564), suggesting an increase in velocity that aligns with the previously observed increase in vertical OKN gain.

Altogether, our data suggest that vestibular-dominant conflict appears to reduce visually dependent oculomotor reflexes, whereas visual-dominant conflict does not impair these capacities.

### Vestibulo-Ocular Reflex suppression

In the VR protocol, horizontal head movements were 79° ± 19° and lasted 671 ms ± 118 ms. In the vertical orientation, movements were 57° ± 14° and lasted 746 ms ± 146 ms. Peak horizontal head velocity was 389°.s⁻¹ ± 154°.s⁻¹, and vertical head peak acceleration was 260°.s⁻¹ ± 79°.s⁻¹. Peak head acceleration was 2135°.s⁻² ± 1233°.s⁻² (horizontal) and 1391°.s⁻² ± 611°.s⁻² (vertical).

In the PF protocol, horizontal head movements were 72° ± 12° and lasted 538 ms ± 75 ms. In the vertical orientation, movements were 53° ± 12° and lasted 572 ms ± 95 ms. Peak horizontal head velocity was 499°.s⁻¹ ± 109°.s⁻¹, and vertical head peak acceleration was 350°.s⁻¹ ± 74°.s⁻¹. Peak head acceleration was 3254°.s⁻² ± 1119°.s⁻² (horizontal) and 2402°.s⁻² ± 776°.s⁻² (vertical).

Peak head velocity remained unchanged post-conflict compared to pre-conflict in the VR, both horizontally (PRE = 386 ± 162, POST = 391 ± 147, F(1,26) = 0.17, p = 0.687) and vertically (Vertical : PRE = 261 ± 80, POST = 259 ± 79, F(1,26) = 0.06, p = 0.803). The PF protocol also showed stable peak head velocity post-flight compared to pre-flight in horizontal (PRE = 488 ± 96, POST = 502 ± 123, F(1,10) = 0.01, p = 0.914) and vertical orientations (PRE = 347 ± 67, POST = 354 ± 83, F(1,11) = 0.04, p = 0.849). Similarly, no significant pre-post differences were found in peak head acceleration, confirming that head movements remained consistent across both time points in VR protocol (Horizontal : PRE = 2136 ± 1288, POST = 2134 ± 1188, F(1,26) = 0.0003, p = 0.987 ; Vertical : PRE = 1409 ± 623, POST = 1372 ± 604, F(1,26) = 0.46, p = 0.505) and PF protocol (Horizontal : PRE = 3096 ± 986, POST = 3360 ± 1229, F(1,10) = 0.12, p = 0.731 ; Vertical : PRE = 2330 ± 697, POST = 2474 ± 857, F(1,11) = 0.14, p = 0.712).

VOR gain significantly decreased after VR exposure in both horizontal (PRE = 0.44 ± 0.15, POST = 0.39 ± 0.13, F(1,26) = 13.2, p = 0.00120, η² = 0.34) and vertical orientations (Vertical : PRE = 0.39 ± 0.1, POST = 0.35 ± 0.09, F(1,26) = 12.2, p = 0.00176, η² = 0.32) (Figure 4C). By contrast, in the PF protocol no significant pre-post effects were observed in either horizontal (PRE = 0.54 ± 0.15, POST = 0.47 ± 0.12, F(1,10) = 2.4, p = 0.151 ; or vertical orientations (PRE = 0.42 ± 0.1, POST = 0.44 ± 0.21, F(1,11) = 0.07, p = 0.799) (Figure 4C). No significant directional effects were observed on VOR gain in either the VR (Horizontal : L-to-R = 0.41 ± 0.14, R-to-L = 0.38 ± 0.15, F(1,26) = 1.9, p = 0.180 ; Vertical : T-to-B = 0.38 ± 0.1, B-to-T = 0.36 ± 0.09, F(1,26) = 0.52, p = 0.477) or PF protocols (Horizontal : L-to-R = 0.53 ± 0.14, R-to-L = 0.54 ± 0.14, F(1,10) = 0.004, p = 0.951 ; Vertical : T-to-B = 0.40 ± 0.17, B-to-T = 0.44 ± 0.16, F(1,11) = 0.37, p = 0.557)(Figure 4B).

Eye Deviation Variability significantly decreased post-VR in the horizontal orientation (PRE = 72.6 ± 33, POST = 63.4 ± 29.4, F(1,26) = 11.4, p = 0.00235, η² = 0.30), with a similar but non-significant trend in the vertical orientation (PRE = 42.1 ± 15.5, POST = 39.4 ± 16.1, F(1,26) = 3.9, p = 0.0601) (Figure 4E). In the PF protocol, EDV tended to decrease in the horizontal orientation (PRE = 108.8 ± 39.3, POST = 88 ± 30, F(1,10) = 3.1, p = 0.106) (Figure 4E), while no trend was observed in the vertical orientation (PRE = 54.5 ± 18.6, POST = 50.3 ± 24.8, F(1,11) = 0.4, p = 0.541) (Figure 4E). In the VR protocol, a directional effect was observed on EDV, with significantly higher values during left-to-right (L-to-R = 64.6 ± 33.3, R-to-L = 62.7 ± 29.6, F(1,26) = 5.1, p = 0.0316, η² = 0.17) and bottom-to-top movements (T-to-B = 35.2 ± 13.5, B-to-T = 46.3 ± 16.0, F(1,26) = 13.9, p < 0.001, η² = 0.35) (Figure 4D). In the PF protocol, a similar directionnal effect was observed on EDV in. vertical direction (T-to-B = 43.6 ± 14.5, B-to-T = 61.2 ± 24.5, F(1,11) = 26, p < 0.001, η² = 0.70) (Figure 4D).

We did not observe any pre-post conflict effects on VOR delay in either the VR protocol (Horizontal : PRE = 183 ± 52, POST = 185 ± 63 ; F(1,26) = 0.1, p = 0.753 ; Vertical : PRE = 224 ± 74, POST = 237 ± 77 ; F(1,26) = 2.7, p = 0.110) or the PF protocol (Horizontal : PRE = 139 ± 36, POST = 134 ± 42 ; F(1,10) = 0.02, p = 0.883 ; Vertical : PRE = 158 ± 38, POST = 157 ± 52 ; F(1,11) = 0.006, p = 0.939).

In conclusion, visually dominant conflict tends to impair vestibulo-driven reflexes, whereas vestibulo-vestibular conflict exhibits more subtle effects.

### Posturography and VR

The 95% area of the center-of-pressure (CoP) displacement was used to assess postural instability, representing the total CoP displacement area during each trial (Figure 5A).

A significant effect of VR exposure was observed, with an increase of 95% area in all three conditions. This increase was significant in the vision deprivation condition (PRE = 78.2 ± 67, POST = 136.9 ± 174.6, W = 86, Z = −2.058, p = 0.020, r = −0.471) (Figure 5B) whereas trend was noted in the fixed vision condition (PRE = 112 ± 91, POST = 191.2 ± 237.1, W = 105, Z = −1.547, p = 0.063, r = −0.354) and in normal vision condition (PRE = 89.6 ± 70.8, POST = 113.9 ± 97.4, W = 124, Z = −1.036, p = 0.156, r = −0.237) (Figure 5B). This suggests that vestibular processing is impaired after VR exposure, an effect that becomes more apparent when visual compensation is unavailable.

In conclusion, absence of visual information might favor postural instability post-VR exposure via potential modifications of vestibulo-proprioceptive integration.

### Sensorimotor performance and motion sickness susceptibility

Correlation analyses were performed to examine the relationships between sensorimotor variables and subjective MS scores, aiming to identify potential predictors of MS. For the VR protocol, we hypothesized that horizontal OKN might correlate with nausea scores, as virtual environments predominantly involve horizontal visual motion. In contrast, since perturbations in the PF protocol were primarily vertical and detected by the vestibular system, we proposed that this orientation would be more relevant for assessing correlations between VOR parameters and nausea scores. In both correlation tests, we examined gain from OKN testing and EDV from VORS testing, as these two variables exhibited longitudinal changes after exposure to either sensory conflict paradigm.

In the VR protocol, baseline horizontal OKN gain showed a significant positive correlation (ρ = 0.387, n = 27, p = 0.046) (Figure 6A). However, vertical baseline EDV didn’t exhibit significant correlations with SSQ score changes during the VR paradigm (Figure 6C).

In contrast, in the PF protocol, a significant negative correlation was found between vertical baseline EDV and SSQ score changes (ρ = −0.697, n = 12, p = 0.012) (Figure 6D). No significant correlations were found between baseline horizontal OKN gain and SSQ score changes in PF protocol (Figure 6B).

In conclusion, visuo-oculomotor sensitivity may be associated with cybersickness, while measures reflecting VORS appear more closely related to SMS susceptibility.

## DISCUSSION

In this paper, we have explored sensory modulations following a brief, single exposure to two distinct types of MS. Our findings provide human evidence of sensory reweighting in low-level behavioral reflexes after exposure to sensory conflicts. We demonstrate that cybersickness and space-sickness have differential effects on visual and vestibular integration. Moreover, our results suggest that MS susceptibility is influenced by an individual’s sensitivity to the sensory cues most involved in detecting displacement.

Firstly, both sensory tests yielded to data consistent with responses reported in the literature. Baseline OKN gain was close to 0.8, with an eye velocity range between 20 and 30°.s^-1^, comparable to findings from similar studies (Kanari, 2017; Watanabe, 1986). For VOR, suppression occurred approximately 100-200 ms after the onset of head acceleration, aligning with prior research (Gauthier & Vercher, 1990). VOR gain immediately preceding suppression was around 0.1-0.4, within the range observed in comparable VOR-eliciting movements (Jell, 1988; Jacobson, 2012). Head peak velocity was approximately 350-500°.s^-1^, while voluntary head movements averaged 80° horizontally and 60° vertically. These parameters were set higher than those typically used in similar studies, making VORS more challenging here (Cullen, 1991).

The nature of the sensory information, either perceived or absent, influences sensory reweighting following exposure to sensory conflicts in our protocols. We initially hypothetized that the incongruent sensory modality would exhibit diminished function after exposure to the respective sensory conflicts. Interestingly, in both types of exposure, the sensory cue considered to be the most reliable appears to be the one that was stimulated. The primary reflexes affected by the vestibular challenge induced by parabolic flight are visually driven reflexes, as evidenced by a decrease in OKN gain post-flight. During parabolic flight, visual cues within the cabin remain relatively stable, whereas the cumulative integration of otolithic inputs corresponds to an altitude change of approximately 8000 feet. Following this significant vestibular stimulation, once the plane lands, it is the optokinetic reflex that is reduced. As a spatially oriented phenomenon, OKN is particularly sensitive to vertical motion, which is notably influenced by the parabolic trajectory experienced by a seated passenger. In this scenario, the otolithic modality is the most stimulated, while the visual information is perceived as unreliable, incongruent, and consequently downweighted. In addition, otolithic detection is consistent with non-vestibular graviceptor inputs, including visceral graviceptors and other sensory systems such as proprioceptive signals (von Gierke & Parker, 1994 ; Mittelstaedt, 1996). These systems collectively contribute to an increased proportion of sensory signals oriented in the direction of the perturbation, which are not aligned with visual inputs explaining why OKN is selectively modulated.

Notably, we anticipated an otolitho-canalar conflict under modified gravitational conditions. In such environments, any head rotation along an axis misaligned with the gravitational field generates a discrepancy between the rotational acceleration detected by the semicircular canals (which remain unaffected by gravity) and the signal from the otolithic sensors. The expected variation in otolithic stimulation differs from what occurs on Earth for a similar semicircular-detected head rotational acceleration. According to the neural mismatch theory, this otolitho-canalar incongruence—relative to the Earth-learned ratio could be a key factor in SMS. Based on this framework, we hypothesized that the reliability of vestibular cues in general would be questioned in such situations, leading to a decrease in vestibular weighting. However, since the measured VOR showed little to no alteration post-flight, this hypothesis is not strongly supported, suggesting that otolitho-canalar integration in modified gravity might not generate neural mismatch. A possible explanation is that otolithic angular detection relies on the ratio between utricular and saccular signals, and this ratio remains unaffected by gravity, which influences both utricular and saccular signals in a similar manner. In this case, it would not generate a conflict with semicircular canal information in this case. However, no studies in the current literature have explicitly investigated or confirmed this hypothesis.

In parabolic flight, the otolithic inputs represent a novel and dynamic perturbation, triggering a reweighting of sensory integration to prioritize different sensory cues. On the other hand, in the VR protocol, the absence of vestibular detection of motion was perceived as dysfunction rather than perturbation. Only vestibular-driven reflexes are altered following exposure to the visual challenge induced by VR, with a reduction in VOR gain observed post-exposure. Additionally, postural stability decreases when vision is deprived, meaning when balance depends solely on proprioceptive and vestibular inputs. It confirms that vestibular performance is impaired following a visual challenge. This lack of accelerometric information coherent with visual perceived motion may have reduced reliance on vestibular inputs. Consequently, sensory conflicts involving inactive inputs appear to diminish their contribution. Our results are in agreement with a proposed common mechanism of multisensorial integration, based on Bayesian inference to reduce perceptual uncertainty (Knill & Pouget, 2004). We observed a reduction in the influence of the lesser-stimulated modality—vestibular on the ground and visual in flight.

Other studies have investigated adaptive integrative processes in humans through repetitive exposure and habituation protocols. Morgan and colleagues (2008) demonstrated that neurons in macaque MSTd modulate the weighting of visual information, increasing reliance on vestibular inputs when visual cues are degraded. Similarly, in mice, a two-week visuo-vestibular conflict led to reduced VOR performance, accompanied by decreased synaptic efficiency in the vestibular nuclei (Idoux, 2018). In humans, a study examining MS reduction through a five-day habituation protocol to visuo-vestibular conflict found that participants experienced shortened vestibular time constants and a decrease in MS symptoms during provocative tests (Dai, 2011). These studies are based on recordings conducted either after long-term habituation or during sensory conflict. Unlike those previous researches, the reweighting observed in our protocols occurred following unique or short repetitive exposure, even though neither visual nor vestibular cues were degraded during the post-test phase. This finding highlights an adaptive mechanism wherein a brief exposure of just 20 minutes was sufficient to induce sensory reweighting. Notably, this reweighting was observed in low-level motor reflexes, contrasting with earlier human studies that primarily used perceptual protocols requiring higher cognitive processing (Alais & Burr, 2004). This suggests that adaptation occurs at an early stage of sensory integration, such as within the vestibular nucleus in the case of the VOR.

The divergence in reflex effects across paradigms raises critical questions, as these reflexes share common anatomical pathways. Multisensory visual-vestibular neurons have been shown to facilitate near-optimal cue integration during self-motion perception (monkeys : Angelaki, 2009). Within the vestibular nuclei, position vestibular pause (PVP) neurons receive direct ipsilateral vestibular inputs and project to contralateral motoneurons, which innervate extraocular muscles, thereby supporting the VOR (monkeys : McCrea, 1987 ; Cullen & Roy, 2004). A direct relationship between PVP interneuron sensitivity and VOR gain has been established (monkeys : Roy & Cullen, 1998). Interestingly, it has been shown that PVP neurons also discharge during OKN and optokinetic after-nystagmus (OKAN) tests, indicating that neuronal mechanisms are shared between VOR and OKN (monkeys : Waespe & Henn, 1977 ; Raphan, 1979 ; Yakushin, 2017). PVP neurons decrease their firing during VORS (monkeys : Roy & Cullen, 1998), likely due to inhibition from the cerebellar flocculus and ventral paraflocculus (monkeys : Lisberger, 1994a, 1994b). The cerebellar flocculus plays a pivotal role in visuo-vestibular adaptation by integrating current sensory inputs from vestibular and non-vestibular input with prior experiences (rabbits : Nagao, 1990 ; monkeys : Stone & Lisberger, 1990 ; mice : Katoh, 1998; Shin, 2011). Based on this framework, our sensory conflict paradigms likely involves discrepancy detection within the cerebellar flocculus, which in turn directly modulates PVP neuron activity. Notably, the downregulation of oculomotor reflexes observed here is not generalized and does not suggest a broad reduction in PVP neuron sensitivity. Instead, it is specific to the source of conflict and the affected sensory modality. This idea is supported by previous researches, demonstrating a 30% decrease in PVP neuron modulation during VORS tasks (involving both visual and vestibular inputs) compared to VOR in darkness (relying solely on vestibular inputs) (monkeys : Cullen & McCrea, 1993 ; Scudder & Fuchs, 1992). Following VR exposure, inhibition of PVP neurons—facilitated by a fixation point— would be reinforced in the VORS task, without altering PVP activity during the OKN task. Conversely, after vestibular challenge during PF, the activation of PVP neurons by visual field motion would be diminished in the OKN task, yet this reduction would not facilitate the downregulation of PVP firing in response to vestibular input during the VORS task.

Given the shared neural pathways and integrative mechanisms between VOR and OKN, it is essential to understand how individual sensory sensitivities influence the detection of discrepancies, multisensory integration processes, and ultimately, susceptibility to sensory conflicts. Our findings reveal a significant relationship between baseline OKN variables and cybersickness severity, with individuals reporting higher nausea scores exhibiting stronger OKN performance metrics. This suggests that heightened visual sensitivity may predispose individuals to more pronounced visuo-vestibular conflicts, aligning with recent findings by Fulvio et al. (2021), who reported a positive correlation between visual cue sensitivity and cybersickness severity in virtual reality. However, in the parabolic flight setting, OKN variables are not significantly correlate with questionnaire score differences, indicating that visual dominance alone may not fully explain MS susceptibility in altered gravitational environments. In contrast, VORS performances, which are not reliable predictors of cybersickness severity, correlate with space sickness severity. In parabolic flight, baseline VORS eye deviation is strongly negatively correlated with SSQ nausea score differences, meaning that participants with better VOR suppression may be more vulnerable to SMS, potentially due to an underlying reduction in vestibular sensitivity. If these individuals naturally rely less on vestibular cues, a further decrease in vestibular efficiency during parabolic flight could amplify their MS symptoms. Similar findings were reported by Thornton (1991) where SMS susceptible individuals displayed higher corrective saccade frequency and less efficient VOR suppression than non-susceptible participants. Collectively, these studies suggest that visual and vestibular sensitivity play a crucial role in MS susceptibility, with the relevant sensory modality depending on the specific type of MS.

## FUNDINGS

This work was supported by CNES (Centre National d’Etudes Spatiales, France) focused on observations with parabolic flights within VP-177 and VP-181 campaigns, and received financial support from the French government in the framework of the University of Bordeaux’s IdEx “Investments for the Future” program / GPR BRAIN_2030.

## DISCLOSURES

No conflicts of interest, financial or otherwise, are declared by the authors.

## AUTHOR CONTRIBUTIONS

*T.B. and E.G.* : Conceived and designed the study, performed experiments, analyzed data, conducted statistical analyses, interpreted results and wrote manuscript.

*E.D.* : Performed experiments

*J.C.* : Contributed to results interpretation and assisted with manuscript revisions

*D.G.* : Assisted in project conception and participant inclusion visit

## Notes

### Competing Interest Statement

The authors have declared no competing interest.

